# A base-line cellular antiviral state is maintained by cGAS and its most frequent naturally occurring variant rs610913

**DOI:** 10.1101/2021.08.24.457532

**Authors:** Julia Kazmierski, Carina Elsner, Katinka Döhner, Shuting Xu, Aurélie Ducroux, Fabian Pott, Jenny Jansen, Christian W. Thorball, Ole Zeymer, Xiaoyi Zhou, Roman Fedorov, Jacques Fellay, Markus W. Löffler, Alexander N. R. Weber, Beate Sodeik, Christine Goffinet

## Abstract

Upon recognition of aberrantly located DNA, the innate immune sensor cGAS activates STING/IRF-3-driven antiviral responses. Here we characterized the ability of a specific variant of the cGAS-encoding gene *MB21D1*, rs610913, to alter cGAS-mediated DNA sensing and viral infection. rs610913 is a frequent G>T polymorphism resulting in a P^261^H exchange in the cGAS protein. Data from the International Collaboration for the Genomics of HIV suggested that rs610913 nominally associates with HIV-1 acquisition *in vivo*. Molecular modeling of cGAS(P^261^H) hinted towards the possibility for an additional binding site for a potential cellular co-factor in cGAS dimers. However, cGAS(WT) or cGAS(P^261^H)-reconstituted THP-1 cGAS KO cells shared steady-state expression of interferon-stimulated genes (ISGs), as opposed to cells expressing the enzymatically inactive cGAS(G^212^A/S^213^A). Accordingly, cGAS(WT) and cGAS(P^261^H) cells were less susceptible to lentiviral transduction and infection with HIV-1, HSV-1, and Chikungunya virus as compared to cGAS KO- or cGAS(G^212^A/S^213^A) cells. Upon DNA challenge, innate immune activation appeared to be mildly reduced upon expression of cGAS(P^261^H) compared to cGAS(WT). Finally, DNA challenge of PBMCs from donors homozygously expressing rs610913 provoked a trend towards a slightly reduced type I IFN response as compared to PBMCs from GG donors. Taken together, the steady-state activity of cGAS maintains a base-line antiviral state rendering cells more refractory to ISG-sensitive viral infections. Even though rs610913 failed to grossly differ phenotypically from the wild-type gene, its expression potentially results in a slightly altered susceptibility to viral infections *in vivo*.

## Introduction

Pattern Recognition Receptors (PRRs) of the innate immune system are crucial for the detection of invading pathogens and required to mount an effective immune response. The cyclic-GMP-AMP-synthase (cGAS) binds to double-stranded DNA in the cytosol and nucleus, followed by its enzymatic activation and the production of the second messenger molecule 2’3’-cyclic GMP-AMP (cGAMP) (1, 2). This small molecule, in turn, binds to the Stimulator of IFN Genes (STING), leading to its activation, phosphorylation and eventually induction of a TANK binding kinase (TBK1)/Interferon regulatory factor 3 (IRF3)-dependent signaling cascade, resulting in the transcription of interferons (IFNs) and IFN-stimulated genes (ISGs), many of them exerting antiviral activity.

cGAS-mediated recognition of invading pathogens serves as a first-line defense mechanism against multiple viruses, which themselves evolved strategies to counteract cGAS-mediated sensing. The genome of invading DNA viruses, such as HSV-1 or KSHV, is recognized in a cGAS-dependent fashion (reviewed in Ref. 3, 4,5). As a consequence, herpes viruses evolved specific antagonists that counteract cGAS/STING-mediated DNA sensing, including HSV-1 pUL41 which selectively targets cGAS mRNA for degradation (6), HSV-1 ICP27 which prevents cGAS phosphorylation (7), or HSV-1-pUL36 which targets STING to proteasomal degradation (8) and therefore interferes with the activation of the crucial transcription factor IRF3. Retroviruses, including HIV-1, evolved a sophisticated replication strategy. Specifically, reverse transcription of their RNA genome into a single DNA intermediate that is destined for integration into a host celĺs chromosome allows retroviruses to largely escape general innate immune activation (9) and cGAS-dependent recognition (10). These observations are in line with studies reporting that innate sensing of HIV-1 infection only occurs upon pharmacological or genetic destabilization of the otherwise nucleic acid-shielding viral capsid (11, 12), and is enhanced in the absence of functional TREX1 expression, that otherwise degrades capsid-escaping and thus cytosolic HIV-1 DNA (13, 14). Interestingly, also RNA viruses have been considered to be inhibited by cGAS-exerted functions, although not mediated through sensing of viral nucleic acids. Rather, cGAS may maintain a basal antiviral state through recognition of self DNA, including endogenous retroelements (15) and/or sensing of DNA released from the nucleus or mitochondria through infection-associated stress induction (16, 17).

Single nucleotide polymorphisms (SNPs) in genes encoding PRRs and downstream adapter molecules modulate infection susceptibility and disease outcome. A remarkable example is the variant of the STING-encoding *TMEM173* gene that contains three non-synonymous SNPs referred to as ‘the HAQ haplotype’. Homozygous expression of this haplotype is predominantly found in East Asian (16.07%) and South American (7.78%) populations (18). It is associated with lower susceptibility to stimulation by cyclic dinucleotides (19) and eventually a severely reduced ability to induce IFN-β expression (19, 20).

Interestingly, among other homozygous SNPs in the *TMEM173* gene, the HAQ haplotype has a higher prevalence in HIV-1 long-term non-progressors, as compared to HIV-1 non-controllers (21).

To date, there is limited knowledge on the role of SNPs in the cGAS-encoding gene *MB21D1*, in particular on implications for DNA sensing and innate immune activation. The most frequent SNP in *MB21D1* is rs610913, a G>T polymorphism that displays a global allele frequency of T = 0.503 (22). The G to T nucleotide exchange results in a single amino acid exchange from histidine (H) to proline (P) at position 261 in the protein sequence. Here, we report structural and functional consequences of the rs610913-encoded P^261^H single amino acid exchange in the cGAS protein in the context of DNA sensing and restriction of viral infections.

## Materials and Methods

### Genome-wide association analysis

Summary statistics for HIV-1 acquisition in the region of *MB21D1* were obtained from genome-wide association analyses (GWAS) previously performed by the International Collaboration for the Genomics of HIV (ICGH) (23). The summary statistics were available on a sub-group basis, with a total of 6 groups matched by geographic origin and genotyping platform, as previously described; Group 1 (The Netherlands, Illumina), Group 2 (France, Illumina), Group 3 (North America, Illumina), Group 4 (French European, Illumina), Group 5 (North American, Affymetrix), and Group 6 (non-Dutch/non-French European, Affymetrix). Association results across groups were combined using a fixed-effects inverse-variance weighted meta-analysis.

### Molecular modeling

The structural model of hcGAS(P^261^H)•dsDNA assembly in the active (ATP-bound) conformational state was created using the ladder-like crystal structure of mouse cGAS in complex with dsDNA (PDB-code: 5N6I) (24) and the structure of the wild type hcGAS•dsDNA•ATP complex (PDB-code: 6CTA) (25). The protein part of the hcGAS•dsDNA•ATP complex was used to generate a homology model of hcGAS(P^261^H) in the active conformational state. The homology model of hcGAS(P^261^H) was superimposed on the mcGAS molecules in the ladder-like assembly. The superposition was performed using the program package Coot (26). The secondary structure matching algorithm (SSM) (27) was used to align the structurally conserved parts of the proteins. The resulting model was subjected to an energy minimization procedure using the program HyperChem (Hypercube, Inc.) with AMBER force field (28) and a distance-dependent dielectric constant. The structural analysis and rendering of Figure 1B and C were performed with the final energy minimized model using the programs COOT and PyMOL (The PyMOL Molecular Graphics System, Version 1.8 Schrödinger, LLC).

**FIGURE 1.**
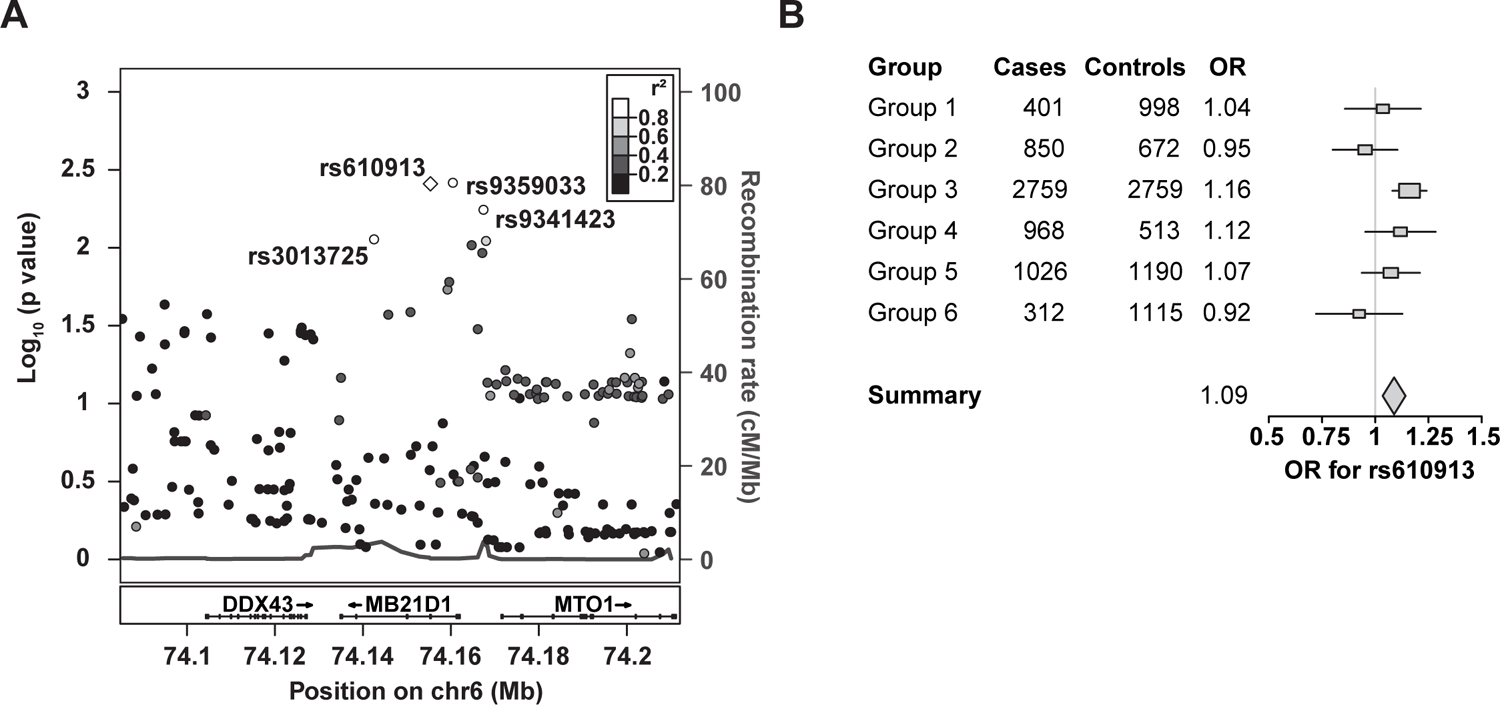
rs610913 may facilitate HIV-1 acquisition *in vivo* (**A**) Regional association plot of the *MB21D1* region, containing all SNPs included in the meta-analysis and their respective p-values. The plot is centered on rs610913 (purple diamond), with red dots indicating SNPs in high linkage disequilibrium (LD) (r^2^>0.8), and green and light blue dots representing SNPs in moderate or low LD. All SNPs in high LD with rs610913 are synonymous SNPs. (**B**) Forest plot of the odds ratios (OR) for rs610913 with 95% confidence intervals across all subgroups and after meta-analysis (diamond) within the ICGH GWAS of HIV-1 acquisition. The number of cases and controls are indicated for each group along with their respective odds ratios.

### Healthy study subjects and blood sample acquisition

Healthy blood donors were recruited at the Interfaculty Institute of Cell Biology, Department of Immunology, University of Tübingen. All healthy blood donors included in this study provided their written informed consent before study participation. Approval for use of their biomaterials was obtained by the respective local ethics committees (approvals 156-2012BO1 and 354-2012BO2), in accordance with the principles laid down in the Declaration of Helsinki as well as applicable laws and regulations.

### Cell lines and primary cells

cGAS KO THP-1 cells (a kind gift from Veit Hornung, Ludwig-Maximilians University, Munich) were cultured in RPMI 1640 medium supplemented with 10% fetal calf serum (FCS), 100 U/ml penicillin, 100 µg/ml streptomycin, 2 mM L-glutamine, 1x MEM non-essential amino acids solution and 1 mM sodium pyruvate (Thermo Fisher Scientific, Waltham, Massachusetts, USA). HEK293T, HEK293T STING-mcherry (1) and BHK-21 cells were maintained in DMEM cell culture medium supplemented with 10% fetal calf serum (FCS), 100 U/ml penicillin, 100 µg/ml streptomycin and 2 mM L-glutamine. HL116 cells (29) were cultured under identical conditions, except for the addition of 1x hypoxanthine-aminopterin-thymidine (HAT) media supplement (Thermo Fisher Scientific, Waltham, Massachusetts, USA). THP-1 cGAS KO cells and HEK293T STING-mcherry cells were reconstituted with individual cGAS-GFP variants by lentiviral transduction and were maintained under 1 µg/ml puromycin selection. After preparation of PBMCs from EDTA-anticoagulated blood by Ficoll-Hypaque centrifugation, cells were stimulated with IL-2 (10 ng/ml) and phytohaemagglutinin (PHA) (1 µg/ml) for 3-4 days, resulting in cultures containing >90% CD3^+^ T-cells. Cells were maintained in RPMI 1640 containing 10% heat-inactivated fetal calf serum, 100 U/ml penicillin, 100 µg/ml streptomycin, 2 mM L-glutamine, 1x MEM non-essential amino acids solution, and 1 mM sodium pyruvate (Thermo Fisher Scientific, Waltham, Massachusetts, USA).

### Generation of lentiviral vector particles and virus stocks

VSV-G-pseudotyped lentiviral vector particles encoding GFP or luciferase were generated by calcium phosphate-based transfection of HEK293T cells with the packaging plasmid pCMV ΔR8.91 (30), the lentiviral transfer plasmids pHR-GFP (31) or pCSII-EF-luciferase (32) and pCMV-VSV-G (33). For the generation of cGAS-transducing lentiviral particles, the transfer plasmids pWPI cGAS(WT)-GFP, pWPI cGAS(G^212^A/S^213^A)-GFP (34), and pWPI cGAS (P^261^H)-GFP were used. pWPI cGAS(P^261^H)-GFP was generated by site-directed mutagenesis (Stratagene California, La Jolla, California, USA), and the correct introduction of the mutation was confirmed by Sanger sequencing. Vector-containing supernatant was collected 40 and 64 hours post-transfection and subjected to ultracentrifugation through a 20% sucrose cushion. To remove residual plasmid DNA, concentrated virus stocks were DNase I-digested twice and stored in aliquots at −80°C. VSV-G-pseudotyped HIV-1 NL4.3 luciferase reporter virus was produced by calcium phosphate-based transfection of HEK293T cells with a HIV-1 NL4.3 ΔEnv ΔVpr luciferase reporter plasmid (35) and a VSV-G-encoding plasmid. Virus-containing supernatants were harvested 60 hours post-transfection and concentrated by ultracentrifugation.

HSV-1 Δ*UL41*N (HSV-1(KOS) UL41NHB) encoding a truncated version of pUL41 was kindly provided by David A. Leib (36). To prepare concentrated stocks, extracellular virions were pelleted from the medium of cells infected at a multiplicity of infection (MOI) of 0.01 PFU/cell for 3 days (37–39). Virus stocks were plaque-titrated on Vero cells (38, 40). To determine the genome/PFU ratio of HSV-1 stocks, we quantified the number of HSV-1 genomes by quantitative PCR as described previously (37, 41).

The CHIKV 181/25 infectious stock (42) expressing a nano-Luciferase fused to the E2 glycoprotein (a kind gift from G. Simmons, Vitalant Research Institute) was produced by *in vitro*-transcription of the full-length, linearized molecular DNA clone into RNA and subsequent RNA electroporation into BHK-21 cells. Virus-containing supernatant was collected three days post electroporation, filtered through membranes of 0.45 µm pores and stored in aliquots at −80°C.

### Flow cytometry

For quantification of cGAS-GFP expression in transduced THP-1 or HEK293T cells, cells were PFA-fixed and GFP positivity was quantified by flow cytometry. HSV-1-challenged HEK293T cells were PBS-washed, PFA-fixed and immunostained for intracellular HSV-1 VP5 using rabbit anti-HSV-1 VP5 (#SY4563) and an appropriate fluorochrome-conjugated secondary antibody in 0.1% Triton in PBS (10, 43). Samples were analyzed on a FACS Lyric device (Becton Dickinson, Franklin Lakes, New Jersey, USA) with BD Suite Software for analysis.

### Immunoblotting

Cell lysates were generated with M-PER Mammalian Protein Extraction Reagent (Thermo Fisher Scientific, Waltham, Massachusetts, USA), run on a 10% SDS-PAGE and transferred onto nitrocellulose using a semi-dry transfer system (Bio-Rad Laboratories, Hercules, California, USA). BSA-blocked membranes were incubated with the primary antibodies mouse-anti human actin (#8226, Abcam, Cambridge, UK), rabbit-anti human cGAS (#15102, Cell Signaling, Danvers Massachusetts, USA), rabbit-anti human IRF3 (#4302, Cell Signaling), rabbit-anti human pIRF3 (#29047, Cell Signaling), rabbit-anti human pSTING (#19781 Cell Signaling), rabbit-anti human pTBK1 (#5483, Cell Signaling), rabbit-anti human STING (#13647S, Cell Signaling), rabbit-anti human TBK1 (#3504, Cell Signaling) or rabbit-anti human TREX1 (#185228, Abcam). Secondary antibodies conjugated to Alexa680/800 fluorescent dyes were used for detection and quantification by Odyssey Infrared Imaging System (LI-COR Biosciences Lincoln, NE, USA).

### Quantitative RT-PCR

Total RNA from cells was extracted using the RNeasy Mini Kit (Qiagen, Hilden, Germany), and residual DNA contaminations were removed with the RNase-free DNase set (Qiagen, Hilden, Germany). Following cDNA synthesis (New England Biolabs, Ipswich, Massachusetts, USA), quantification of relative mRNA levels was performed using Taq-Man PCR technology (Thermo Fisher Scientific, Waltham, Massachusetts, USA) with primer-probe kits (Applied Biosystems, Waltham, Massachusetts, USA) for following genes: *ACTB* (Hs03023943_g1), *ARL16* (Hs01586770_g1), *BST2* (Hs00171632_m1), *cGAS* (Hs00403553_m1), *HAUS7* (Hs00213860_m1), *IFIT1* (Hs01911452_s1), *IFN-β* (Hs01077958_s1), *IRF3* (Hs01547283_m1), *LYAR* (Hs00215132_m1), *MX2* (Hs01550814_m1), *RPL30* (Hs00265497_m1), *RPS11* (Hs06642555_g1), *STING* (Hs00736958_m1), *TCP1* (Hs01053946_g1), *TREX1* (Hs03989617_s1), *TRMT10C* (Hs01933516_s1), *YBX1* (Hs00358903_g1).

Relative mRNA levels were determined in multiplex reactions using the ΔΔCt method with human *RNASEP* mRNA as an internal reference. Each sample was analyzed in technical triplicates and with parallel controls omitting reverse transcriptase. Assays were performed on an OneStep Plus machine (Applied Biosystems, Waltham, Massachusetts, USA) or a LightCycler 480 II (Roche, Basel, Switzerland). Data analysis was performed using Applied Biosystems Step One Software (Version 2.3) or LightCycler 480 Software (Version 1.5).

### RNA sequencing

Total RNA extraction from cells and DNase treatment were performed with the RNeasy Mini kit and RNase-free DNase set (Qiagen, Hilden, Germany). The quality and integrity of total RNA were verified on an Agilent Technologies 2100 Bioanalyzer (Agilent Technologies, Waldbronn, Germany). The RNA sequencing libraries were generated using TruSeq Stranded mRNA Library Prep Kits (Illumina, San Diego, California, USA) following the manufactureŕs protocol. The libraries were sequenced on Illumina HiSeq4000 (paired-end run 2 x 75 bp) with an average of 9 x 10^7^ reads per RNA sample. Data generated from individual samples were mapped separately against the hg38 human reference genome. Gene expression was calculated for individual transcripts as reads per kilobase per million bases mapped (RPKM). All transcriptomic analyses were performed using Geneious Prime Version 2020.0.4 (Biomatters, New Zealand). Differentially expressed genes (DEGs) were identified by calculating fold changes in expression, genes were considered to be expressed significantly differently if their expression was increased by at least a factor of two with a p-value of < 0.05. Gene ontology enrichment analyses were performed using the Panther overrepresentation test (http://geneontology.org/), Homo sapiens reference list, and GO biological process complete annotation data set. p-values were corrected using a false discovery rates (Ashburner et al., 2000; The Gene Ontology Consortium, 2019).

### Electroporation

10 million THP-1 cells and PBMCs were electroporated (140 V, 1000 µF) in serum-free RPMI in the presence of endotoxin-free plasmid DNA (12 µg DNA, or indicated quantities), 4 µg cGAMP (Invivogen, San Diego, Californien, USA), 4 µg c-di-UMP (Invivogen, San Diego, Californien, USA) or mock-electroporated using a Gene Pulser Xcell Electroporation instrument (Bio-Rad Laboratories, Hercules, California, USA) and 0.2 cm cuvettes

### HL116 cell-based detection of bioactive IFNs

Culture supernatants of individual cell lines were titrated on HL116 cells that express the luciferase gene under the control of the IFN-inducible 6-16 promoter (29). Six hours post-inoculation, cells were PBS-washed and luciferase expression was determined using Cell Culture Lysis Buffer (Promega, Madison, Wisconsin, USA) and the Luciferase Assay System (Promega, Madison, Wisconsin, USA). The concentration of IFN was quantified using an IFN-α2a (Roferon) standard curve.

### Protein Purification

The full-length human cGAS(WT) and cGAS(P^261^H) proteins were expressed in *E.coli* BL21(DE3). The expression was induced by 0.5 M isopropyl-β-D-thiogalactoside and incubated at 18°C for 18 h. After centrifugation at 5,000 x g for 15 min, pellets were resuspended in 20 ml PBS and centrifuged again at 5000 x g for 15 min. The cells were flash-frozen and stored at −80°C until purification. For purification, pellets were thawed and resuspended in a buffer containing 300 mM NaCl, 50 mM Na_3_PO_4_, 10 mM imidazole, pH 7.5 with complete protease inhibitor cocktail (Roche, Basel, Switzerland) and lysed by sonification for 2 min, with 4 min breaks after eachminute of sonification to prevent overheating of the lysate. DNase I was added to remove a possible impurity caused by the cellular DNA bound to cGAS. After 30 minutes of incubation at room temperature, the sample was centrifuged at 40,000 x g for 1 h. The supernatant was filtered using a 0.45 µm syringe filter and loaded onto a 5 ml Protino Ni-NTA column (Macherey-Nagel, Düren, Germany). The column was washed until UV280 had reached a steady value and eluted with 500 mM imidazole, 150 mM NaCl, 50 mM Na_3_PO_4_, pH 7.5. The pooled fractions were diluted with low salt buffer (50 mM NaCl, 20 mM Tris, pH 7.5) to prevent protein aggregation caused by a high salt concentration of the elution buffer. The diluted eluate was then loaded onto a 1 ml HiTrap Heparin HP column (GE Healthcare, Chalfont St Giles, UK). After loading, the column was washed until UV280 reached a steady value before elution with an increasing salt gradient buffer: 50 mM – 2 M NaCl, 20 mM Tris, pH 7.5. The elution was concentrated by centrifugation with 30,000 MWCO Vivaspin Hydrosart (Sartorius, Göttingen, Germany) and, if needed, diluted with low salt buffer to the final protein concentration used for the *in vitro* activity assay. The purified protein was flash-frozen in liquid nitrogen and stored at −80°C.

### *In vitro* cGAS activity assay

2 µM recombinant human cGAS was incubated for 2h at 37°C with the substrates 0.5 mM ATP and 0.5 mM GTP, in the presence of 1 ng/µl dsDNA fragments NoLimits (Thermo Fisher Scientific, Waltham, Massachusetts, USA) of 1, 4 or 6 kb length, in a buffer containing 120.25 mM MnCl_2_, 20 mM NaCl, 2.5 mM MgCl_2_, 8 mM Tris-HCl, pH 7.5 in a total volume of 200 µl. Following incubation, samples were inactivated at 95°C for 20 min. Samples were centrifuged at 14500 x rpm for 15 min at 4°C, and the supernatant was diluted 1:100 with H_2_O for HPLC measurement with the API 4000 LC-MS/MC System (Sciex, Framingham, Massachusetts, USA) for 2’3’cGAMP quantification. The gradient started with 100% buffer A (3/97 (v/v) MeOH/H_2_O, 50 mM NH_4_Ac, 0.1% HAc) and reached 50% buffer A, 50% buffer B (97/3 (v/v) MeOH/H_2_O, 50 mM NH_4_Ac, 0.1% HAc) after 5 min. The sample was run over a ZORBAX Eclipse XDB-C18 1.8 µm, 50 × 4.6 mm (Agilent Technologies, Waldbronn, Germany) column. Measurements and data generation were controlled by Analyst software (version 1.5.2., SCIEX). Calibration was conducted with 10 µl of synthetic derived 2’3’cGAMP mixed with 800 µl extraction reagent (2/2/1 [v/v/v] methanol, acetonitrile and water mixture) and 300 µl extraction reagent (25 ng/mL tenofovir in extraction reagent) with tenofovir as the internal standard.

### Infection and transduction assays

30 min prior to lentiviral transduction, cells were left untreated or treated with Efavirenz (EFV; 100 nM). Inhibitor treatment was maintained during the subsequent virus inoculation. Transduction and virus infection assays were performed by spin-inoculation for 60 min at 32°C. Following spinoculation, cells were incubated at 37°C, 5% CO_2_, and individual wells were harvested at indicated time points.

### Luciferase assay

Luciferase expression of cells challenged with VSV-G lentiviral vectors or VSV-G/HIV-1 NL4.3 was quantified 72 hours post-transduction. Cells were washed with PBS, lysed using Cell Culture Lysis Buffer (Promega, Madison, Wisconsin, USA) and luciferase activity was quantified using the Luciferase Assay System (Promega, Madison, Wisconsin, USA). To detect nanoluciferase expression in supernatants from CHIKV-infected cells, the Nano-Glo Luciferase Assay System (Promega, Madison, Wisconsin, USA) was used according to manufacturer’s instruction.

### LPS and poly(I:C) treatment

IL-2/PHA-activated PBMCs were treated with 40 ng/ml LPS or 20 µg/ml poly(I:C) as previously described (46, 47).

### Reagents and inhibitors

Fragmented dsDNA (NoLimits 100 bp DNA Fragment) for *in vitro* experiments were obtained from Thermo Fisher. Lipopolysaccharide (LPS) and poly (I:C) were purchased from Sigma-Aldrich (St. Louis, Missouri, USA). Human IFN-α2a (Roferon) was purchased from Roche. 2’-3’-cGAMP and c-di-UMP were purchased from Invivogen. EFV was purchased from Bristol-Myers Squibb.

### Genotyping of PBMCs for rs610913 (Tübingen)

200 µl freshly drawn EDTA-anticoagulated venous whole blood (S-Monocette K3 EDTA, Sarstedt, Hümbrecht, Germany) was subjected to DNA isolation using the QIAamp DNA Blood Mini Kit on a Qiacube (both from Qiagen) instrument following the manufacturer’s instructions. For genotyping an *MB21D1* rs610913-specific Taqman primer set (Thermo Fisher, C_937459_20), diluted 20x, were used with 20 ng genomic DNA and the appropriate amounts of TaqMan Universal MasterMix II (Thermo Fisher) in a 10 µl reaction volume run in triplicate wells of a 96 well MicroAmp plate run on a QuantStudio 7 qPCR cycler (Thermo Fisher) and QuantStudio Real-Time PCR Software v1.3.

### Data Availability

RNA-seq datasets will be deposited at the NCBI GEO database.

### Data presentation and statistical analysis

If not otherwise stated, bars and symbols show the arithmetic mean and error bars the S.E.M of the indicated number of individual experiments. Statistical significance was calculated by performing paired Student’s t-test using GraphPad Prism 8. *p* values <0.05-0.01 were considered significant (*) and <0.01 very significant (**) or not significant (p-value ≥0.05; n.s.).

## Results

### rs610913 may facilitate HIV-1 acquisition *in vivo*

In the human population, the coding region of the cGAS-encoding gene *MB21D1* harbors several non-synonymous SNPs of different frequencies (**Table 1**). The allele frequencies vary substantially across populations, with rs610913 being the most frequent non-synonymous SNP (**Table 1**). We searched for SNPs in *MB21D1* displaying an association with HIV-1 acquisition using summary statistics covering 6,334 infected cases and 7,247 controls of European ancestry (23) (**Fig. 1A**). Interestingly, we detected a nominal, however not genome-wide significant over-representation of rs610913 (p=0.004) in HIV-1-infected individuals as compared to the uninfected control cohort, suggesting that this SNP may associate with and/or facilitate HIV-1 acquisition. Analysing rs610913 in more detail across the six included subgroups, indicated that the signal was primarily arising from Group 3, a group consisting of North American individuals and enriched for HIV controllers (23) (**Fig. 1B**). Given its high global frequency and its potential role in HIV-1 acquisition, we embarked on a functional study of rs610913.

**Table 1.** Allele frequency of most abundant, non-synonymous SNPs in the cGAS-encoding gene *MB21D1* Shown are the respective single nucleotide exchange, resulting amino acid substitution and relative allele frequencies of the reference and alternative alleles in the African, European and global populations.

| SNP | Alleles | Amino Acids | Total |  | African |  | Europe |  |
| --- | --- | --- | --- | --- | --- | --- | --- | --- |
|  |  |  | Reference | Alternative | Reference | Alternative | Reference | Alternative |
| *rs9352000* | G>T | T35N | 15,65 | 84,35 | 14,37 | 85,63 | 16,28 | 83,72 |
| *rs610913* | G>T | P261H | 49,72 | 50,28 | 63,09 | 36,91 | 35,59 | 64,41 |
| *rs35629782* | G>T | A48E | 94,98 | 5,02 | 98,44 | 5,51 | 94,49 | 5,51 |
| *rs141133909* | C>T | G101R | 97,95 | 2,05 | 99,38 | 0,62 | 97,79 | 2,21 |
| *rs145259959* | A>G | L239P | 99,80 | 0,20 | 99,98 | 0,02 | 99,78 | 0,22 |
| *rs138984002* | T>A | Y483F | 99,96 | 0,04 | 99,74 | 0,26 | 100,00 | 0,00 |
| *rs114473784* | C>T | E422K | 99,96 | 0,04 | 98,73 | 1,27 | 100,00 | 0,00 |
| *rs141390590* | G>T | F433L | 99,98 | 0,03 | 100,00 | 0,00 | 99,97 | 0,03 |
| *rs146116825* | G>C | S393C | 99,98 | 0,02 | 99,90 | 0,10 | 100,00 | 0,00 |

### Structural modeling of cGAS(P^261^H) reveals amino acid position at “head-to-head” interface of the cGAS ladder-like assembly

To investigate the structural impact of cGAS(P^261^H) mutation, we built a molecular model of the hcGAS(P^261^H)•dsDNA assembly in the active (ATP-bound) conformational state. The overall structure of this assembly is based on the experimental ladder-like cGAS•dsDNA crystallographic model obtained for the mouse enzyme (mcGAS) (24). The positions of mcGAS molecules in the ladder-like assembly were substituted by the homology model of the hcGAS(P^261^H) mutant based on the structure of the wild type hcGAS•dsDNA•ATP complex (PDB-code: 6CTA)(25). The model of hcGAS P^261^H•dsDNA•ATP ladder-like assembly was optimized, and the geometry of the resulting model (**Fig. 2**) appeared to be very close to the original mcGAS•dsDNA assembly due to the high structural and sequence similarity (r.m.s.d. 1.0 Å, sequence identity 70%) between the human and mouse enzymes. In the hcGAS(P^261^H)•dsDNA•ATP ladder-like structure, the H^261^ residue is located far away from the active site in the deep cleft created by the “head-to-head” interface between the two hcGAS monomers bound to the dsDNA (**Fig. 2**). Another H^261^ residue from the neighboring “head-to-head” hcGAS(P^261^H) molecule is located at the bottom of the same site. Together two imidazole rings of the H^261^ residues create a positively charged surface at the bottom of the “head-to-head” hcGAS P^261^H cleft (**Fig. 2D**). The distance between the two H^261^ residues in the cleft is rather large (∼11 Å), which makes direct interaction between them unlikely. The distances between H261 and the two dsDNA molecules (9.3 Å and 15.6 Å) also do not allow direct contact (**Fig. 2C**). At the same time, the side chain of H^261^ makes two new hydrogen bonds with the side chains of S^201^ and E^259^ of the same monomer, which is not possible for the proline side chain of P^261^ in the WT enzyme (**Fig. 2C**). These hydrogen bonds could provide additional stabilization of cGAS(P^261^H) monomers in the “head-to-head” cleft and may contribute to the overall stability of the cGAS•dsDNA assembly. Since S^201^, E^259^, and H^261^ residues are located in a solvent-accessible area, the free energy of their interaction may be expected to be diminished by the solvent effects. Thus, the modelling analysis, indicates that cGAS P^261^H is not expected to cause a significant discrepancy in enzymatic activity compared to the WT enzyme, although the overall stability of the multimeric complex with DNA could be affected slightly.

**FIGURE 2.**
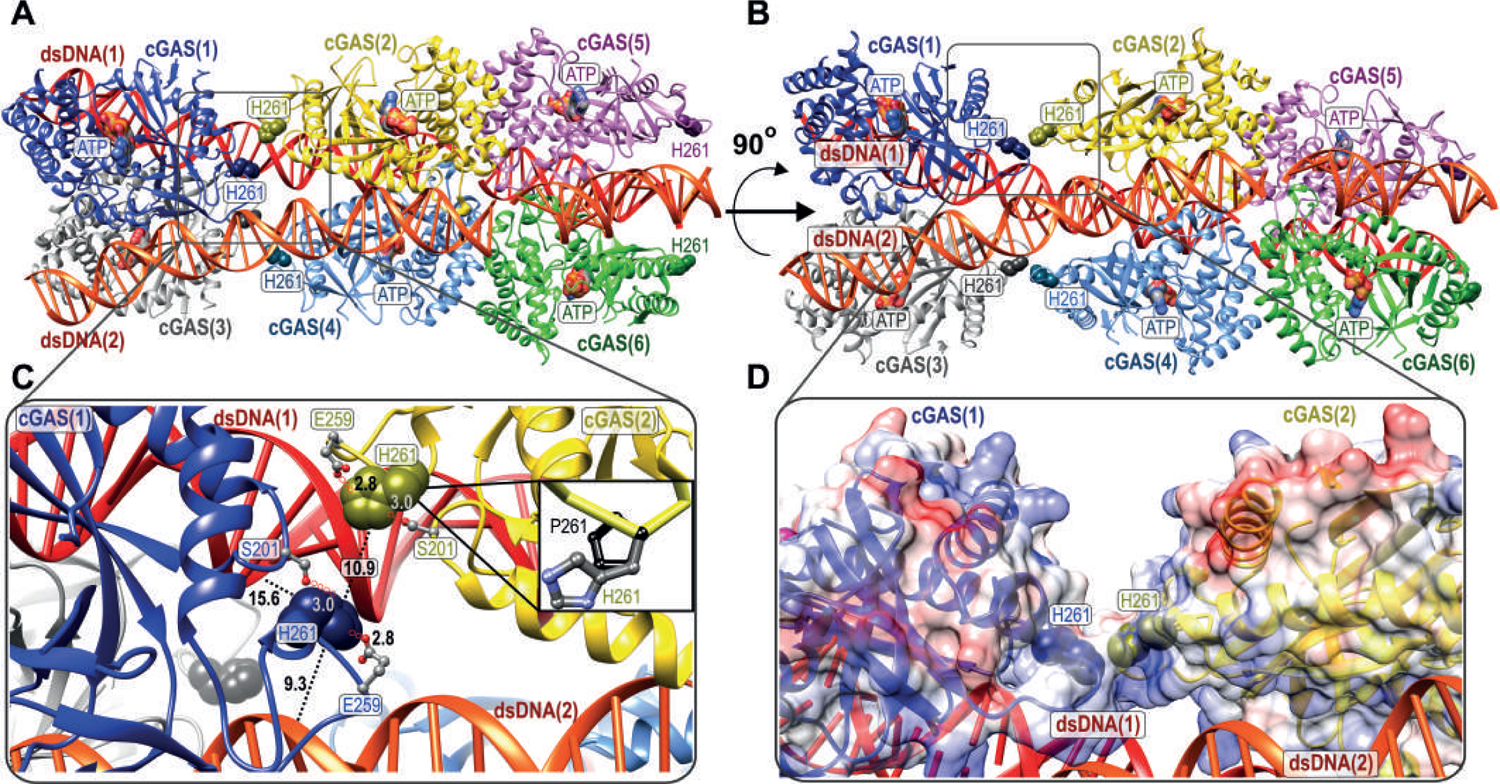
Structural model of cGAS(P^261^H) reveals amino acid position at “head-to-head” interface of the cGAS-ladder-like assembly (**A**) and (**B**) The structural model of human cGAS(P^261^H)•dsDNA oligomeric assembly created using the ladder-like crystal structure of mouse cGAS in complex with dsDNA (PDB-ID: 5N6I) and the structure of the hcGAS(WT)•dsDNA•ATP complex (PDB-ID: 6CTA) as starting coordinates. The cGAS(P^261^H) monomers are shown in blue, yellow, magenta, grey, cyan, and green. The two dsDNA molecules are shown in orange and red. The residues H^261^ in the cGAS(P^261^H) monomers are represented with a Corey-Pauling-Koltun (CPK) model with the corresponding colors. The ATP binding sites are indicated using molecular surface representation. (C) Detailed view of H^261^ localization. The closest distances between H^261^ residues and the dsDNA molecules are shown with dotted black lines. The hydrogen bonds between H^261^, S^201^, and E^259^ are traced with red circles. The close-up panel in the black box shows the comparison of H^261^ and P^261^ side-chain structures. (D) Molecular surface representation of the “head-to-head” interface cleft between the two cGAS(P^261^H) monomers bound to the dsDNA molecules. The semitransparent surface is colored according to the molecular electrostatic potential with positive, negative, and neutral charges represented by the blue, red, and white colors, respectively. The residues H261 are shown using the CPK model representation.

### Catalytically active cGAS modulates base-line levels of *IFIT1*, *MX2* and *IFNB1* mRNA expression

In order to address functional consequences that may result from the proline-to-histidine exchange encoded by rs610913, we stably expressed individual cGAS-GFP variants in THP-1 cGAS KO cells by lentiviral transduction, including cGAS(WT)-GFP, catalytically inactive cGAS(G^212^A, S^213^A)-GFP (48) and cGAS(P^261^H)-GFP. Assessment of GFP expression by flow cytometry and of cGAS expression by immunoblotting confirmed similar expression levels of the transgenes in individual cell lines, as opposed to mock-transduced THP-1 cGAS KO cells (**Fig. 3A**). Other key components of the cGAS signaling cascade, such as STING, IRF3, and TREX1, were expressed at similar levels in the four cell lines (**Fig. 3B**), indicating that abrogation of cGAS expression or of its catalytic activity did not affect mRNA or protein quantities of gene products involved in this pathway. Interestingly, basal expression of *IFIT1*, *MX2*, and *IFNB1* mRNA was clearly reduced in cells devoid of cGAS expression and in cells expressing catalytically inactive cGAS, when compared to cells expressing either WT cGAS or cGAS(P^261^H) (**Fig. 3C**).

**FIGURE 3.**
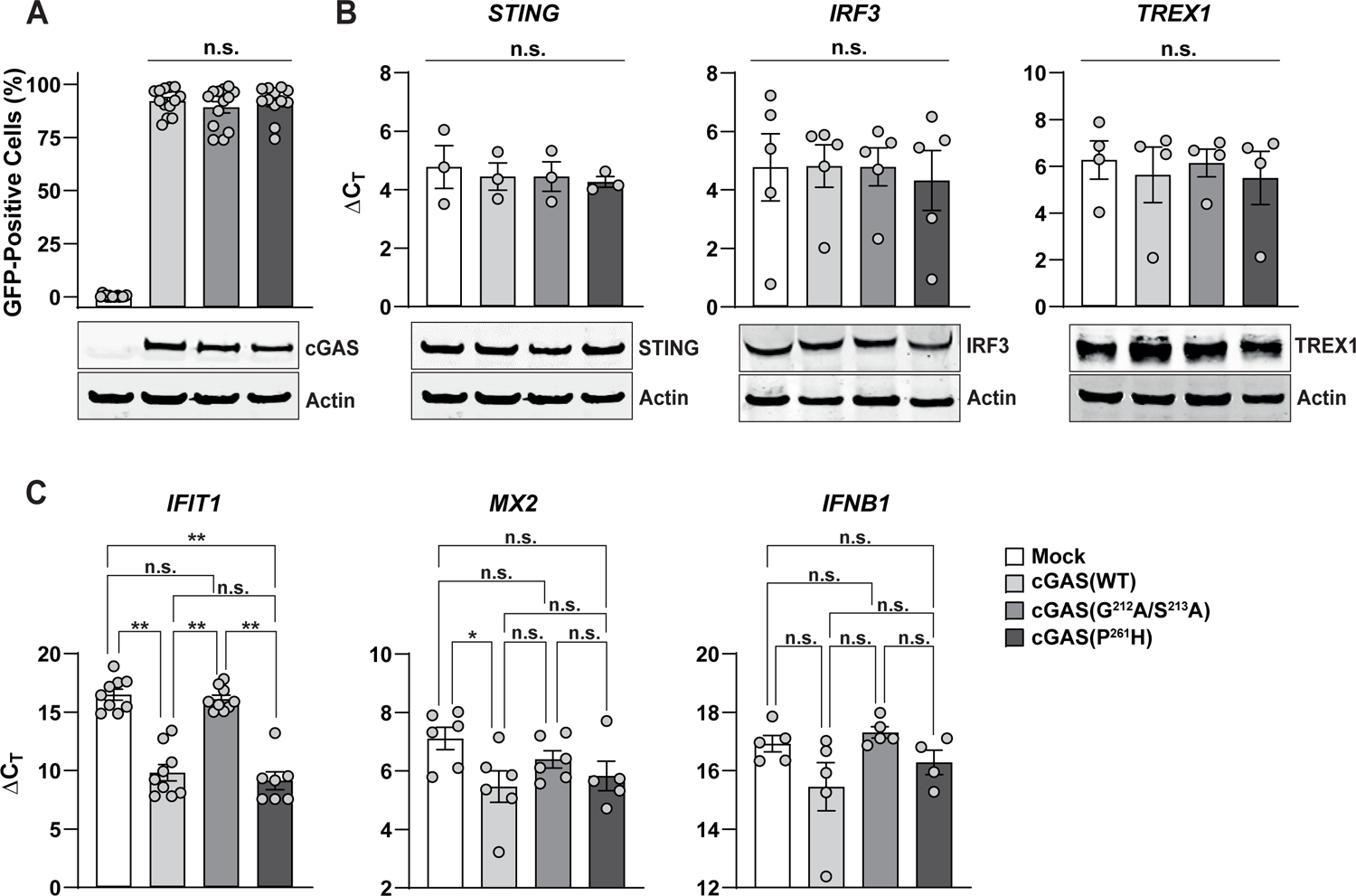
Catalytically active cGAS modulates base-line levels of *IFIT1*, *MX2* and *IFNB1* mRNA expression (**A-C**) THP-1 cGAS KO cells were stably transduced with indicated GFP-cGAS variants and analysed for: (A) Percentage of GFP-positive cells and steady-state cGAS protein expression by flow cytometry and immunoblotting, respectively. (B) Relative expression of *STING*, *IRF3* and *TREX1* mRNA by quantitative RT-PCR and immunoblotting of indicated proteins. (C) Relative *IFIT1*, *MX2* and *IFNB1* mRNA expression levels by quantitative RT-PCR. Error bars indicate S.E.M from ≥ three independent experiments. Immunoblots shown are representative blots of two or more.

### Expression of functional cGAS induces global transcriptomic alterations in THP-1 cells

To explore transcriptional profiles associating with expression of individual functional and non-functional cGAS variants, we subjected total RNA of indicated THP-1 cells to sequencing. Plotting of all RPKM values >0.5 revealed a high overall correlation in the gene expression profile between the individual samples (**Fig. 4A**). cGAS(WT) and cGAS(P^261^H) cells (**Fig. 4A, top panel**) shared a similar expression profile. Comparison of cGAS(WT) or cGAS(P^261^H) the catalytically inactive cGAS(G^212^A/S^213^A) revealed a set of 77 and 115 genes significantly upregulated exclusively in the context of the functional cGAS variants, respectively, suggesting that expression of those genes requires cGAS base-level activity (**Fig. 4B, middle and bottom panels**). Interestingly, gene ontology analysis revealed that the genes whose expression was overrepresented in cGAS(WT) and cGAS(P^261^H)-expressing cells, as compared to cGAS (G^212^A/S^213^A) cells, joined common gene sets, including cellular defense mechanisms to invading pathogens (GO:0009615 Response to Virus; GO:0051607 Defense Response to Virus) or type I IFN signaling (GO:0034340 Response to Type I IFN; GO:0060337 Type I IFN Signaling Pathway) (**Fig. 4C, middle and bottom panels**). In accordance, the fifty most differentially expressed genes (DEGs) among all significant DEGs in cGAS(WT) compared to cGAS(G^212^A/S^213^A) samples represented mostly ISGs (43 ISGs out of 50 DEGs), such as *IFI44L*, *IFI27* and *MX1* or important components of the type I IFN signaling axis, such as *STAT1* and *IRF7* (**Fig. 4D**). In line with previous experiments, known components or modulators of the cGAS/STING signaling axis were equally expressed throughout all cell lines, independent of functional cGAS expression (**Sup. Fig. 1A**).

**FIGURE 4.**
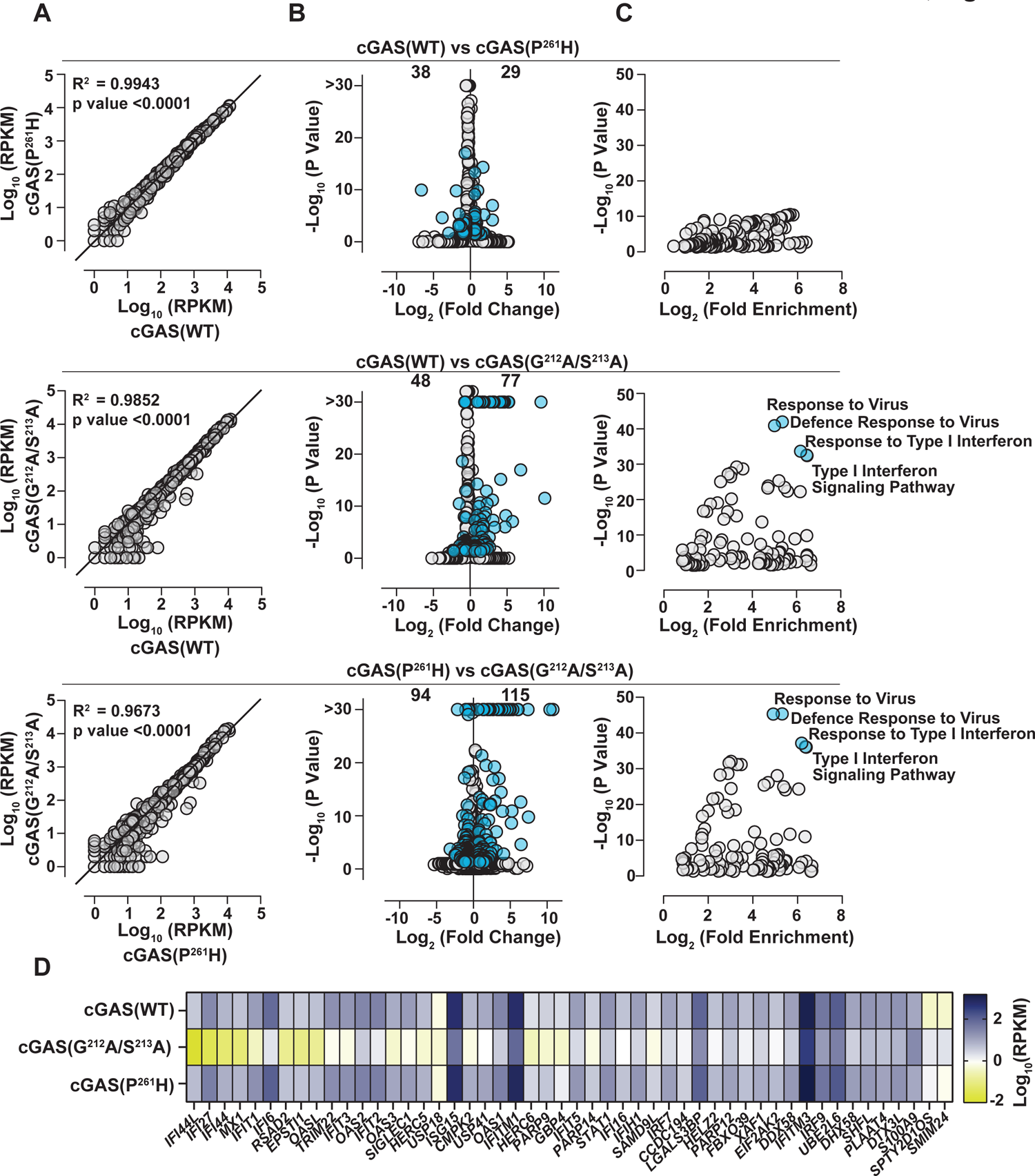
Expression of functional cGAS induces global transcriptomic alterations in THP-1 cells (**A-D**) Bulk RNA sequencing analysis was conducted with total RNA extracted from THP-1 cGAS KO cells stably expressing indicated cGAS variants. (A) Plot of all raw RPKM values > 0.5 of the indicated samples. (B) Identification of differentially expressed genes in cGAS (WT) vs cGAS (P^261^H), cGAS (WT) vs cGAS (G^212^A/S^213^A) or cGAS (P^261^H) vs cGAS (G^212^A/S^213^A) samples by plotting the gene expression fold change and *p*-values of all differentially expressed genes. Genes with a *p*-value < 0.05 and more than two-fold change are highlighted in blue. (C) Gene ontology analysis of genes significantly upregulated in cGAS (P^261^H) vs cGAS (WT), cGAS (WT) vs cGAS (G^212^A/S^213^A) or cGAS (P^261^H) vs cGAS (G^212^A/S^213^A) expressing cells. (D) Heatmap showing the RPKM values of the 50 DEGs of highest absolute fold change of all statistically significant (*p* < 0,05) DEGs in cGAS(WT) vs cGAS(G^212^A/S^213^A) samples. Genes are ranked based on their absolute fold change.

Although the overall transcriptome of cGAS(WT) and cGAS(P^261^H) expressing THP-1 cells appeared very homogenous (**Fig. 4A, top panel**), 67 genes were DEGs which reached statistical significance (**Fig. 4B, top panel**). These genes, however, displayed low or moderate expression fold changes and *p*-values. In addition, gene ontology enrichment analysis revealed enrichment of gene sets with only moderate *p*-values and divergent functions (**Fig. 4C, top panel**), indicating that expression of cGAS(P^261^H) does not severely modulate the cellular transcriptome. We selected ten candidate genes based on fold change and statistical significance (**Sup. Fig. 1B**) and evaluated their expression by Q-RT-PCR (**Sup. Fig. 1 C)**, aiming at challenging the findings obtained with RNA sequencing. In line with the rather subtle differences in the transcriptomes of cGAS(WT) and cGAS(P^261^H)-expressing cells, analysis of several independent samples by Q-RT-PCR confirmed only *TCP-1* out of the ten tested candidate genes as a true DEG whose expression is specifically increased in the context of cGAS(P^261^H) expression, thus displaying lower mRNA levels in cGAS(WT)-expressing cells. In summary, the transcriptomic data provide further evidence for a base-line antiviral immunity in cells expressing functional cGAS(WT) or cGAS(P^261^H), and both cGAS variants control an overall highly similar cellular transcriptome. In contrast, the absence of cGAS expression or expression of a functionally inactive cGAS mutant decreased the antiviral state of the cell as reflected by lower expression of genes related to virus defense and the type I IFN response.

### cGAS(WT) and cGAS(P^261^H) expression reduces susceptibility to lentiviral transduction in the absence of transduction-provoked innate immune responses

Since rs610913 may associate with increased probability of HIV-1 infection *in vivo*, we next investigated whether expression of cGAS(P^261^H) renders cells more susceptible to infection by HIV-1 and other viruses. Specifically, we challenged THP-1 cGAS KO cells reconstituted with cGAS(WT), cGAS(G^212^A/S^213^A) or cGAS(P^261^H) with VSV-G-pseudotyped lentiviral particles or HIV-1 NL4.3 and monitored the transduction efficiency. Interestingly, cells devoid of cGAS expression or expressing the catalytically inactive mutant displayed higher susceptibility to lentiviral transduction as compared to cGAS(WT) or cGAS(P^261^H)-expressing counterparts (**Fig. 5A-B**). However, cGAS(WT) and cGAS(P^261^H)-expressing cells shared identical susceptibility to lentiviral transduction.

**FIGURE 5.**
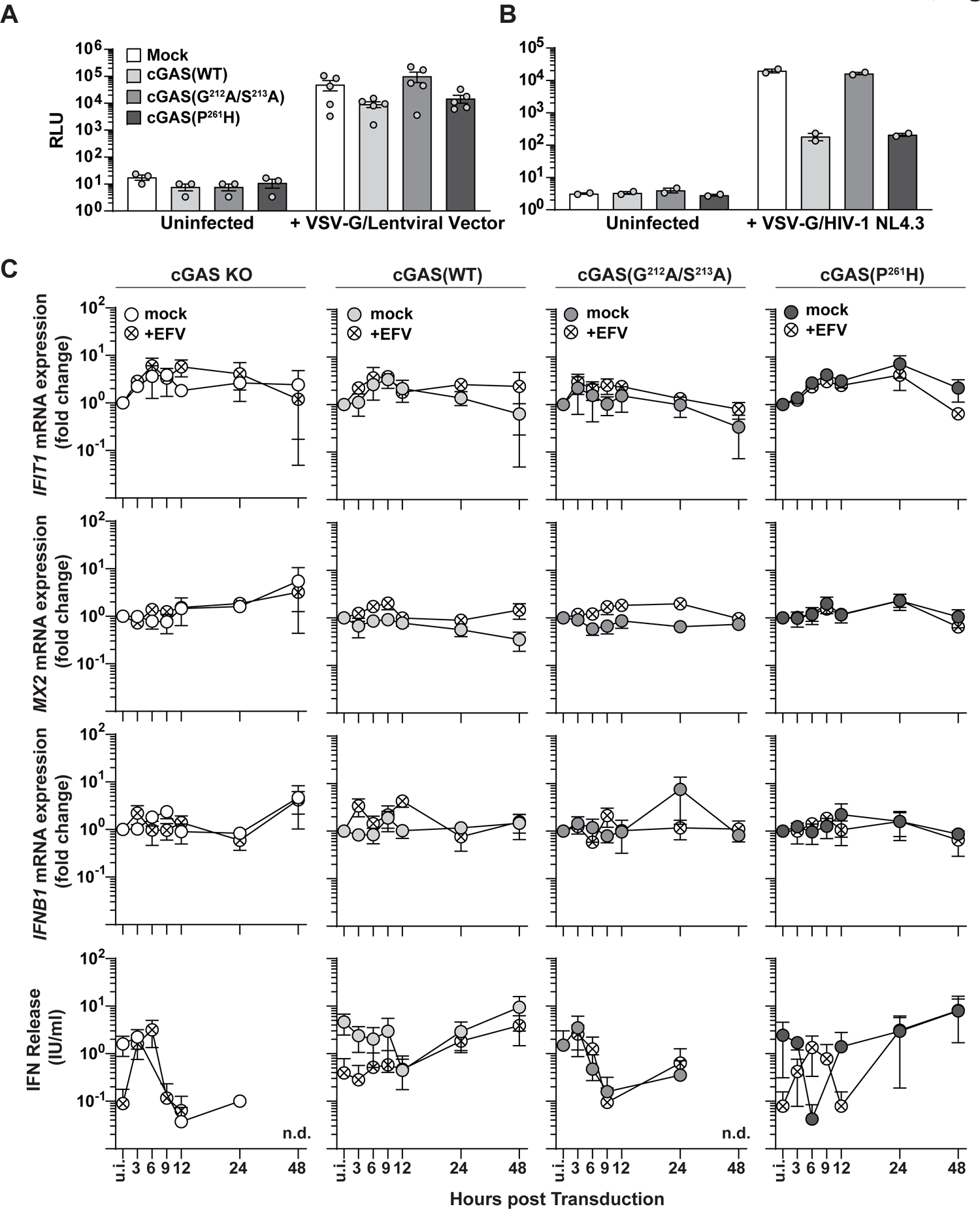
cGAS(WT) and cGAS(P^261^H) expression reduces susceptibility to lentiviral transduction in the absence of transduction-provoked innate immune responses (A) THP-1 cells were transduced with VSV-G-pseudotyped lentiviral vectors and analyzed for luciferase reporter expression 72 hours post transduction (n = 5). (B) THP-1 cells were infected with VSV-G-pseudotyped HIV-1 NL4.3 luciferase and luciferase reporter expression was analyzed 72 hours post transduction (n = 2). (C) Indicated THP-1 cells were transduced with VSV-G-pseudotyped lentiviral vectors in the presence or absence of the reverse transcriptase inhibitor Efavirenz (EFV). Shown is *IFIT1*, *MX2* and *IFNB* mRNA expression and the release of bioactive IFNs in cell culture supernatants at the indicated time points (n = 3).

Importantly, transduction of cells with ablated cGAS expression or expressing individual cGAS variants did neither induce expression of *IFIT1*, *MX2* and *IFNB* mRNAs, nor secretion of bioactive IFN into the culture supernatant in an EFV-sensitive fashion (**Fig. 5C**). Lentiviral transduction triggered induction of *IFIT1* mRNA expression to a maximum of 2.9 to 7.1-fold in all four cell lines, irrespective of their cGAS expression status or EFV treatment. These results are consistent with absence of cGAS-mediated responses to lentiviral infection reported by others (9, 11) and us (10), suggesting that detectable differences in the susceptibility of our cell lines to transduction are linked to different antiviral states.

### Base-line antiviral state mediated by cGAS(WT) and cGAS(P^261^H) expression renders cells less susceptible to HSV-1 and CHIKV infection

To explore the role of cGAS in the context of infection with other viruses displaying individual replication strategies and genomic architectures, we reconstituted HEK293T, that lack detectable cGAS and STING expression, and HEK293T mCherry-STING cells (1) with individual cGAS variants (**Sup Fig. 2 A-B**). In line with results obtained in THP-1 cells, expression of STING, IRF3, and TREX1 was not affected by complementation of cGAS expression in HEK293T cells (**Sup Fig. 2 C-D**). Reconstitution with both cGAS and STING expression restored the cGAS-dependent base-level induction of *IFIT1*, *MX2* and *IFNB1* expression in HEK293T cells, indicating the intactness of the remaining signaling pathway in HEK293T cells (**Sup Fig. 2E**). Based on these observations, we considered cGAS/STING-expressing HEK293T cells as a suitable model to monitor cGAS-dependent restriction of viral infections. As a prototypic DNA virus, we used an HSV-1 strain that encodes a truncated version of pUL41, a well-characterized cGAS antagonist (6, 36). Cells expressing cGAS(WT) or cGAS(P^261^H) were less susceptible to infection with HSV-1 Δ*UL41*N. Their rate of HSV-1 Vp5-positive cells scored to 46.7% and 43.8%, respectively, as opposed to 61.7% in cGAS(G^212^A/S^213^A)-expressing cells and 56.8% in cGAS-negative cells (**Fig. 6A**). Strikingly, the same pattern was observed in the context of Chikungunya virus (CHIKV) strain 181/25, an RNA virus (**Fig. 6B**). Here, cGAS(WT) or cGAS(P^261^H)-expressing cells displayed luciferase reporter expression of 5.180 and 5.177 RLU, respectively, as compared to cGAS(G^212^A/S^213^A)-expressing cells that yielded a mean value of 16.760 RLU. Together, these data support the idea that cGAS maintains a base-line antiviral milieu that acts in a broad manner against invading viral pathogens. Conclusively, beyond sensing viral DNA intermediates or stress-induced host DNA released from intracellular compartments in the during an ongoing infection, cGAS expression and steady-state activity may maintain a static antiviral state that represents a hurdle for viral infections that are sensitive to the cGAS-controlled antiviral ISG program.

**FIGURE 6.**
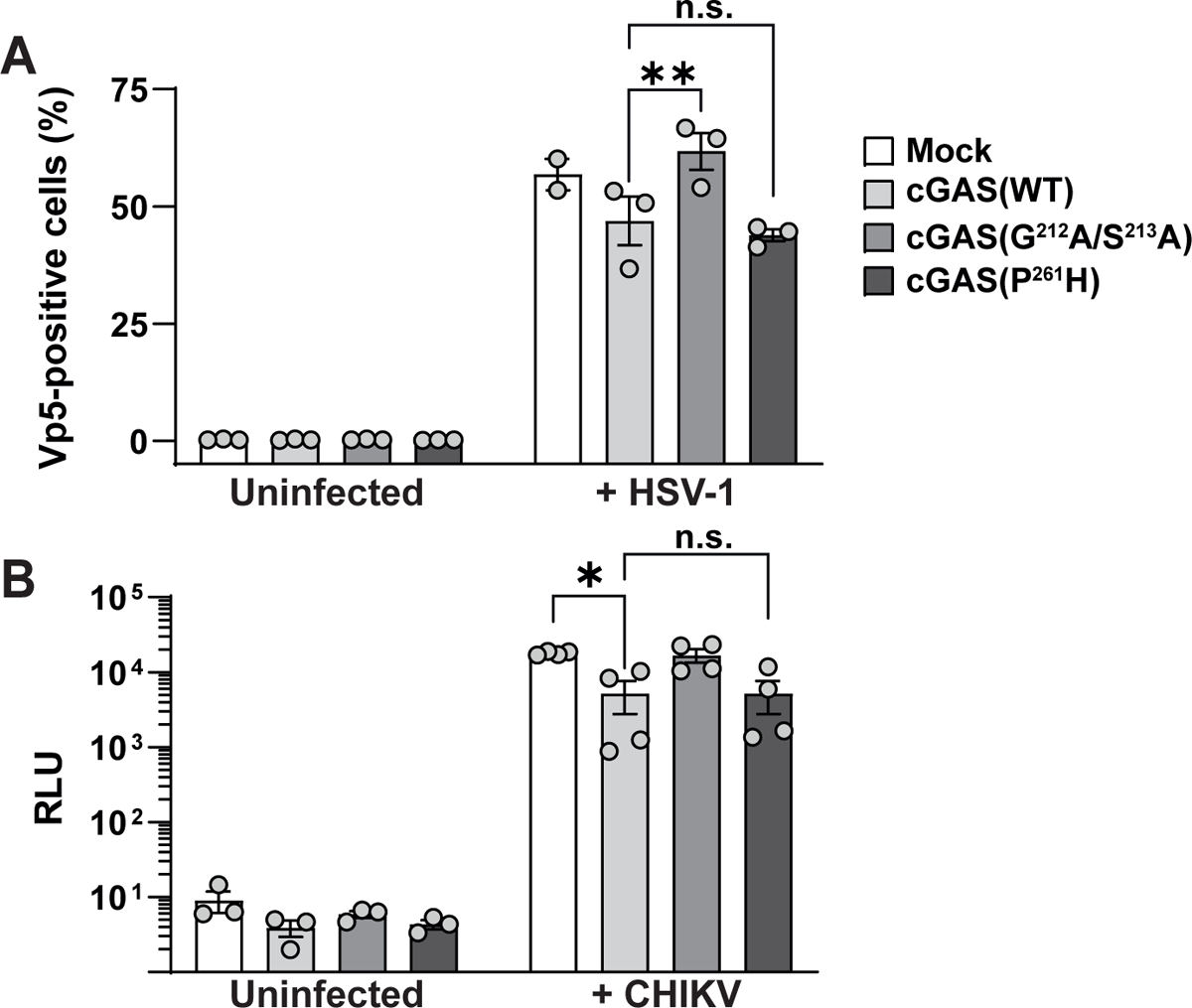
Base-line antiviral state mediated by cGAS(WT) and cGAS(P^261^H) expression renders cells less susceptible to HSV-1 and CHIKV infection (A) 293T mCherry-STING cells stably expressing cGAS variants were challenged with HSV-1 Δ*UL41*N followed by quantification of intracellular HSV-1 Vp5 protein expression by flow cytometry. (B) 293T mCherry-STING cells stably expressing cGAS variants were infected with CHIKV 181/25 luciferase reporter strain followed by luciferase detection 48 hours post infection. Error bars indicate S.E.M. from ≥ 3 individual experiments.

### cGAS(P^261^H) may display a slightly reduced DNA sensing ability

We next investigated the functionality of cGAS-mediated DNA-sensing and induction of the type I IFN response of THP-1 cells expressing cGAS(P^261^H) as compared to cGAS(WT) by quantifying the type I IFN response provoked upon electroporation with endotoxin-free plasmid DNA. Of note, human cGAS is efficiently activated upon binding to long dsDNA, as opposed to binding to short DNA fragments (24)(25). Electroporation of plasmid DNA resulted in the release of bioactive IFN at concentrations of 11,158 IU/ml and 33,652 IU/ml into supernatants of cells expressing cGAS(WT) or cGAS(P^261^H), respectively (**Fig. 7A**). In contrast, cGAS KO cells and cells expressing the inactive cGAS(G^212^A/S^213^A) mutant barely responded to plasmid DNA challenge and displayed responses that did not exceed the levels of mock-electroporated cells. Electroporation with the STING agonist cGAMP, but not the control cyclic dinucleotide c-di-UMP, induced release of similar levels of bioactive IFNs in all tested cell lines, indicating the intactness of the STING signaling axis (**Fig. 7A**). Similarly, phosphorylation of STING, TBK1, and IRF3 upon plasmid DNA challenge was detectable as early as 0.5 and 1 hour post plasmid DNA challenge in cells expressing cGAS(WT) and cGAS(P^261^H), whereas lysates from both THP-1 cGAS KO cells and THP-1 cells expressing cGAS (G^212^A/S^213^A) scored negative in this assay, as expected (**Fig. 7B**). While the quality and kinetics of the type I IFN response upon challenge with a fixed plasmid copy number did not reveal gross differences between cGAS(WT) and cGAS(P^261^H), titration of plasmid DNA uncovered a slightly inferior ability of cGAS(P^261^H) over cGAS(WT) to induce *IFIT1* mRNA expression (**Fig. 7C**), but not release of bioactive IFNs in the cell culture supernatant (**Fig. 7D**). To unravel potentially different inherent catalytic activities of cGAS(WT) and cGAS(P^261^H), both proteins were expressed in *E. coli*, purified and incubated with dsDNA fragments of 1, 4 or 6 kb length in the presence of ATP and GTP. Both proteins presented similar *in vitro* enzymatic activities as judged by 2’-3-cGAMP quantification by LC-MS/MC, with a trend towards higher cGAMP production by cGAS(P^261^H) as compared to cGAS(WT) (**Fig. 7E**). For both cGAS variants, the enzymatic activity increased with augmenting dsDNA length, in accordance with other reports on cGAS(WT) (24, 49). In summary, while our *in vitro* data seem to suggest a slightly inferior *in vitro* catalytic activity of cGAS(WT) as compared to cGAS(P^261^H), our functional data in cells indicate a slightly superior sensitivity of the WT protein to DNA that manifests at suboptimal DNA quantities.

**FIGURE 7.**
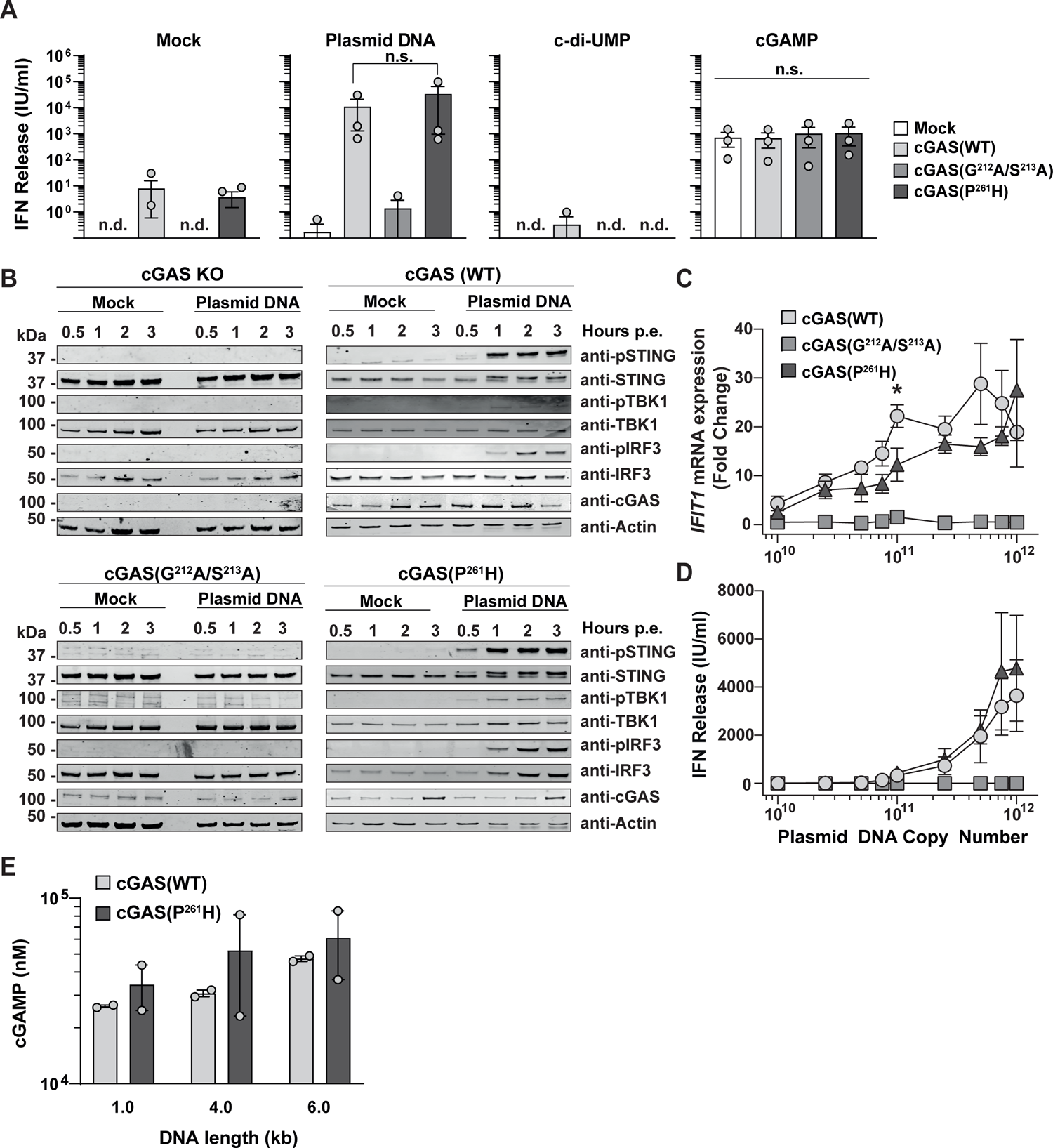
cGAS(P^261^H) may display a slightly reduced DNA sensing ability in THP-1 cells (A) Indicated THP-1 cells were electroporated with plasmid DNA (12 µg), c-di-UMP (6 µg), cGAMP (6 µg) or mock-electroporated. Shown is the release of bioactive IFNs in cell culture supernatants. (B) Immunoblotting of indicated total and phosphorylated proteins were performed using lysates from indicated THP-1 cells electroporated with plasmid DNA (12 µg) or mock-electroporated and collected at indicated time points post challenge. One representative blot of two is shown. (C) *IFIT1* mRNA expression of indicated THP-1 cells electroporated with increasing amounts of plasmid DNA. (D) Release of bioactive IFNs in cell culture supernatant indicated THP-1 cells electroporated with increasing amounts of plasmid DNA. (E) The *in vitro* activity of purified cGAS (WT) and cGAS(P^261^H) proteins in the presence of dsDNA fragments of various lengths (1, 4, and 6 kb) is shown. The *in vitro* activity was measured in terms of 2’3’cGAMP production by cGAS incubated with its substrates ATP and GTP. Error bars indicate S.E.M of two experiments performed with two individual protein purifications. If not otherwise stated error bars indicate S.E.M. from three individual experiments. n.d. = not detectable; p.e. = post electroporation.

### rs610913 homozygosity results in a trend towards a lower cell-intrinsic response to plasmid DNA, but not to LPS and poly(I:C) challenge

According to data from the 1000 genomes project (22), the SNP rs610913 displays an allele frequency of 35.6 to 63.1% in humans. PBMCs from a cohort of healthy individuals were isolated and stratified upon genotyping of corresponding while blood. Steady-state *IFIT1* mRNA expression levels were similar in PBMCs from individuals homozygous for the WT variant and individuals homozygous for rs610913 (**Fig. 8A**). *IFIT1* mRNA expression was slightly, but non-significantly, increased in the rs610913 SNP group compared to the cGAS(WT) group after both LPS and poly(I:C) challenge (**Fig. 8B**). In contrast, plasmid DNA challenge revealed a slightly decreased *IFIT1* mRNA expression in the rs610913 SNP group compared to the cGAS(WT) group. These data point towards the possibility of a slightly impaired DNA sensing ability in the context of the cGAS(P^261^H)-encoding rs610913 gene variant.

**FIGURE 8.**
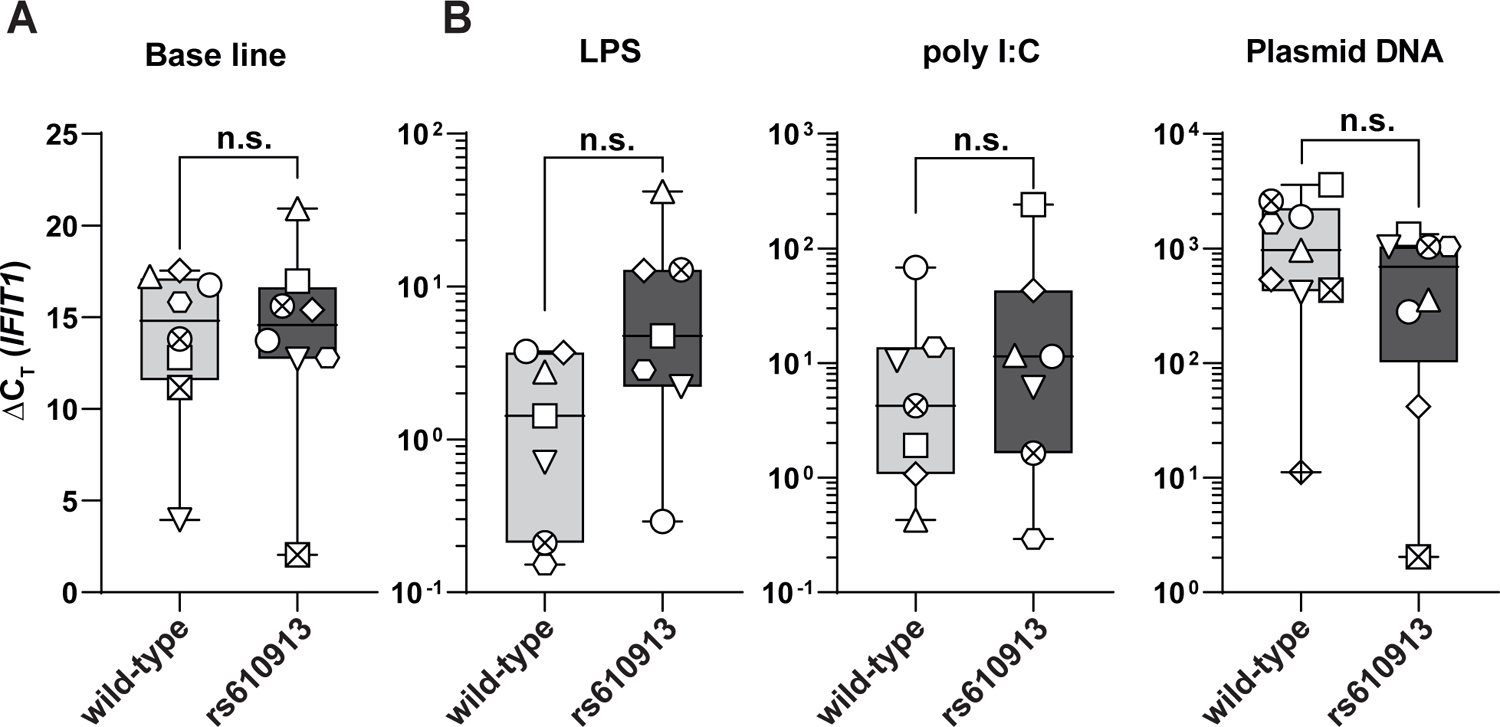
rs610913 homozygosity results in a trend towards a lower cell-intrinsic response to plasmid DNA, but not to LPS and poly(I:C) challenge IL-2/PHA-stimulated PBMCs from healthy donors with indicated genotype were analyzed in respect to: (A) Basal level *IFIT1* mRNA expression of IL-2/PHA-stimulated PBMCs. (B) *IFIT1* mRNA expression in IL-2/PHA-stimulated PBMCs upon poly(I:C), LPS or plasmid DNA challenge. The symbols indicate individual donors. Error bars display S.E.M.

## Discussion

This study aimed at characterizing the impact of a single amino acid exchange, proline-to-histidine at position 261 of the cGAS protein. The SNP rs610913 encoding for cGAS(P^261^H) attracted our attention because of its high allele frequency. With the exception of the protective deleting polymorphism in the *CCR5* gene (*CCR5Δ32*), little genetic contribution on HIV acquisition was identified in previous genome-wide association studies (23, 50). The mild apparent overrepresentation of the rs610913 allele in HIV-1-positive individuals prompted us to hypothesize that it associates with a higher risk of HIV-1 acquisition. Of note, rs610913 appeared to be enriched in BCG-vaccinated healthy controls compared to TB-positive vaccinated individuals, suggesting an association of rs610913 with BCG vaccine-mediated protection to TB infection (51). Given the lack of additional data on rs610913, we aimed at evaluating its role *in vitro*, *in cellulo* and *ex vivo*.

Interestingly, we first detected a strong association of cGAS expression and maintenance of base-level innate immunity, in the absence of infection or external stimuli. Both the expression of individual tested ISGs, and the global transcriptomic profile shifted significantly towards an antiviral state in THP-1 cGAS KO cells upon restoration of cGAS expression. This observation was corroborated in HEK293T cells equipped with both cGAS and STING expression. Also, components of the IFN signaling cascade, such as *STAT1* and *IRF7*, were expressed at higher levels in the presence of functional cGAS, potentially allowing bystander cells to mount more rapid paracrine responses. This observation is reminiscent of our previous findings in mouse CD4^+^ T-cells, which equally displayed a cGAS-dependent ISG expression profile in the absence of exogenous stimuli (10). Work by other groups linked the cGAS-mediated priming of innate immunity to the release and sensing of mitochondrial DNA as response to cellular stress (52, 53), a pathway that can be triggered by both DNA or RNA virus infection (16, 52, 54) and to the base-line sensing of endogenous retroviruses. It is therefore conceivable that cGAS-mediated activity may not only target viruses with dsDNA genomes or DNA intermediates, but also RNA viruses. Along this line, several RNA viruses indeed evolved strategies to actively counteract the cGAS/STING signaling axis (17, 55).

However, comparison of cells expressing cGAS(P^261^H) and cGAS(WT) protein failed to reveal pronounced differences in their ability to maintain a base-line innate immunity in any experimental system we studied, suggesting a similar efficiency of cGAS(P^261^H) enzymatic function, at least at steady-state conditions. At base-line levels, we identified a differential regulation of *TCP-1*, which encodes for a molecular chaperone that is part of the TRiC complex (56). Upon challenge with high amounts of plasmid DNA, cells expressing either cGAS(WT) or cGAS(P^261^H) supported a robust release of similar concentrations of bioactive IFNs, as opposed to cells expressing the non-functional cGAS mutant and cGAS KO cells. Also, kinetics of phosphorylation of STING, TBK1 and IRF3 were similar in cGAS(P^261^H) and cGAS (WT)-expressing cells. In the context of challenge with suboptimal DNA quantities, induction of *IFIT1* mRNA expression was significantly reduced in cGAS(P^261^H)-expressing cells compared to cGAS(WT) cells. Likewise, PBMCs from homozygous rs610913 carriers displayed a trend towards reacting at lower magnitudes to DNA challenge than cells from homozygous WT allele carriers. These data point towards a possibly reduced DNA binding affinity of cGAS(P^261^H) or differential requirement of cGAS cofactor interaction. The latter idea is supported by the results of our molecular modeling attempts, which hinted towards the possibility of a potential additional co-factor binding site in the cGAS(P^261^H) protein. The topology and the surface charge distribution of the “head-to-head” hcGAS(P^261^H) cleft containing the two H^261^ residues create a favorable binding site for a potential cellular co-factor that might increase the stability of the hcGAS P^261^H•dsDNA ladder-like assembly *in vivo*. This stability increase would contribute to the nucleation-cooperativity-based mechanism of cGAS (24) and enhanced enzymatic activity. The latter is supported by the observation that a slightly higher *in vitro* catalytic activity of cGAS(P^261^H) can be attributed to the additional stabilization of the “head-to-head” area by the two hydrogen bonds between H^261^ and the side-chains of S^201^ and E^259^ of the same monomer, which are absent in the WT enzyme. Besides, the presence of two histidine residues in the cleft makes this site more suitable for specific recognition and high-affinity binding compared to proline. Intriguingly, a previous report suggested a loss of helix and glycosylation of the mutated cGAS(P^261^H) protein, and a better capacity for binding interactions (51).

Individual expression of cGAS(WT) or cGAS(P^261^H) conferred a decreased susceptibility to VSV-G-pseudotyped lentiviral vector-mediated and HIV-1 transduction. This inhibition occurred in the absence of detectable induction of an innate immune response upon transduction. These data support the idea that cGAS-induced base-line antiviral state of the cells, rather than cGAS-mediated detection of viral DNA intermediates or infection-triggered release of mtDNA, is responsible for lower transduction efficiencies. Of note, the absence of innate immune activation upon HIV-1 infection has been linked to the intactness of viral capsids that permit capsid uncoating closely tied to nuclear pores or in proximity to integration sites within the nucleus, thereby preventing exposure of HIV-1 RT products to cytosolic DNA sensors (11, 12, 57, 58). In line with our working model, cGAS expression reduced susceptibility also to infection with HSV-1 and CHIKV, an RNA virus. Schoggins and colleagues proposed cGAS-mediated inhibition of RNA virus infection through exerting an IRF3-dependent but STAT-independent mechanism (15). Alphaviruses including CHIKV are sensitive to STING/IRF3-mediated restriction of infection (59, 60). In contrast, herpesvirus infection can be accompanied by accidental leakage of viral DNA into the cytosol, allowing cGAS-mediated recognition of the viral nucleic acids and subsequent type I IFN responses (61, 62). Although we detected a protective role of functional cGAS in our experiments, we failed to establish a specific phenotype of cGAS(WT) compared to the cGAS(P^261^H) variant, indicating that both proteinś expression establishes a cellular antiviral state that sufficiently restricts infection by diverse viruses.

In conclusion, we demonstrate the overall intact functionality of rs610913 SNP-encoded cGAS(P^261^H). This protein, similarly to cGAS(WT) mounts an efficient IFN response upon sensing of dsDNA and decreases susceptibility to infection by different viruses by maintenance of a cGAS-dependent, base-line expression of multiple antiviral factors.

## Acknowledgments

We thank Sandra Pelligrini, Veit Hornung, and Jens Bohne for the kind gift of the HL116 cell line, THP-1 cGAS KO cells, and HEK293T-mcherry-STING, and HEK293T cells, respectively. We thank Oya Cingöz for providing the plasmid HIV-1 NL4.3 ΔEnv ΔVpr luciferase. We thank Victor Tarabykin for granting access to the Step One Plus Real-Time PCR System at Charité Universitätsmedizin. We thank the Genomics platform of the Berlin Institute of Health for NGS. We thank Rune Hartmann and Andreas Holleufer for their help with the *in vitro* activity experiments. We are very grateful to Dietmar Manstein, Rune Hartmann, and Karl-Peter Hopfner for many fruitful discussions. We thank Thomas Pietschmann and Christian Drosten for their constant support. We thank the HIV Reagent Program for providing essential reagents. We thank Sabine Dickhöfer for genotyping support and technical assistance. We thank all study subjects and their families, as well as voluntary healthy blood donors for participating in the study. J.K. is supported by the Center of Infection Biology and Immunity (ZIBI). This work was supported by a postdoctoral fellowship from the Foundation Ernst & Margarete Wagemann to A.D., by funding from German Research Foundation (Deutsche Forschungsgemeinschaft, DFG) to C.G. (Collaborative Research Centre SFB900 “Microbial Persistence and its Control”, Project number 158989968, project C8 and Priority Programm 1923 “Innate Sensing and Restriction of Retroviruses”, GO2153/4 grant) and to B.S. (SFB900 158989968; project C2; EXC2155 RESIST 390874280; So 403/6-1), by funding from Boehringer Ingelheim Foundation (Exploration Grant) to C.G., funding of the Helmholtz Center for Infection Research (HZI) and of Berlin Institute of Health (BIH) to C.G. R.F., X.Z and O.Z. were supported by German Research Foundation grant FE 1510/2-1 and EXC 2155 “RESIST” – Project ID 39087428. ANRW was supported by the Else-Kröner-Fresenius Stiftung, the University Hospital Tübingen, the University of Tübingen, the DFG Clusters of Excellence “iFIT – Image-Guided and Functionally Instructed Tumor Therapies” (EXC 2180, also to MWL) and “CMFI – Controlling Microbes to Fight Infection (EXC 2124). Gefördert durch die Deutsche Forschungsgemeinschaft (DFG) im Rahmen der Exzellenzstrategie des Bundes und der Länder - EXC 2180 and EXC 2124.

## Author Contributions

JK, CE, SX, CG designed research.

JK, CE, KD, SX, AD, FP, JJ, OZ, XZ, RF performed research. MWL and ANRW were involved in sample acquisition

JK, CE, SX, AD, FP, CWB, OZ, XZ, RF, JF, ANRW, BS, CG analyzed data.

CE, KD, CWB, RF, ANRW, BS contributed to writing the manuscript. JK and CG wrote the paper.

## Nonstandard Abbreviations

BCG: Bacille Calmette-Guérin; Vaccine against Tuberculosis

cGAMP: cyclic Guanosine Monophosphate-Adenosine Monophosphate cGAS – Cyclic GMP-AMP Synthase

CHIKV: Chikungunya Virus

DEG: Differentially Expressed Gene

EFV: Efavirenz

IFIT1: Interferon-Induced Protein with Tetratricopeptide Repeats 1

IFN: Interferon

ISG: Interferon-Stimulated Gene

IRF3/7: Interferon Regulatory Factor 3/7

KSHV: Kaposi’s Sarcoma-Associated Herpesvirus

MX2: MX Dynamin Like GTPase 2

PBMC: Peripheral Blood Mononuclear Cell

PHA: Phytohaemagglutinin

PRR: Pattern Recognition Receptor

RLU: Relative Light Unit

RPKM: Reads per Kilobase of Transcript

RT: Reverse Transcriptase

SNP: Single-Nucleotide Polymorphism

STAT1: Signal Transducer and Activator of Transcription 1

STING: Stimulator of Interferon Genes

TBK1: TANK-binding Kinase 1

TREX-1: Three Prime Repair Exonuclease 1

VSV-G: Vesicular Stomatitis Virus Glycoprotein

---

**SUPPLEMENTAL FIGURE 1.**
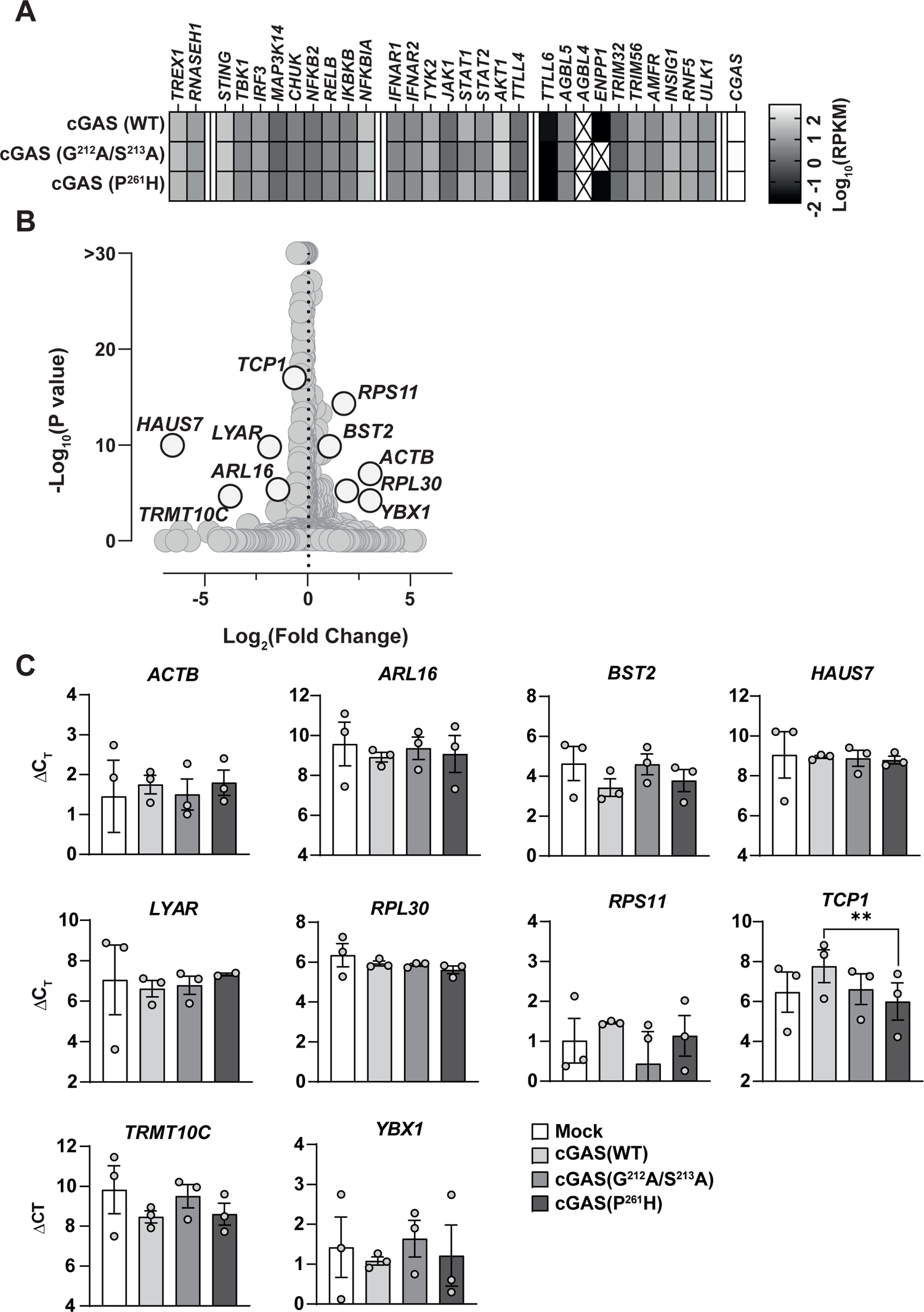
Transcriptomic Analysis of THP-1 cells stably expressing cGAS variants (A) Heatmap showing the raw RPKM values of genes involved in the cGAS/STING signaling axis. (B) Plot of differential expressed genes in cGAS(WT) vs cGAS(P^261^H) samples. Genes that were verified by RT-Q-PCR are highlighted in red. (C) C_T_ values of selected genes normalized to RNASEP expression tested by RT-Q-PCR.

**SUPPLEMENTAL FIGURE 2.**
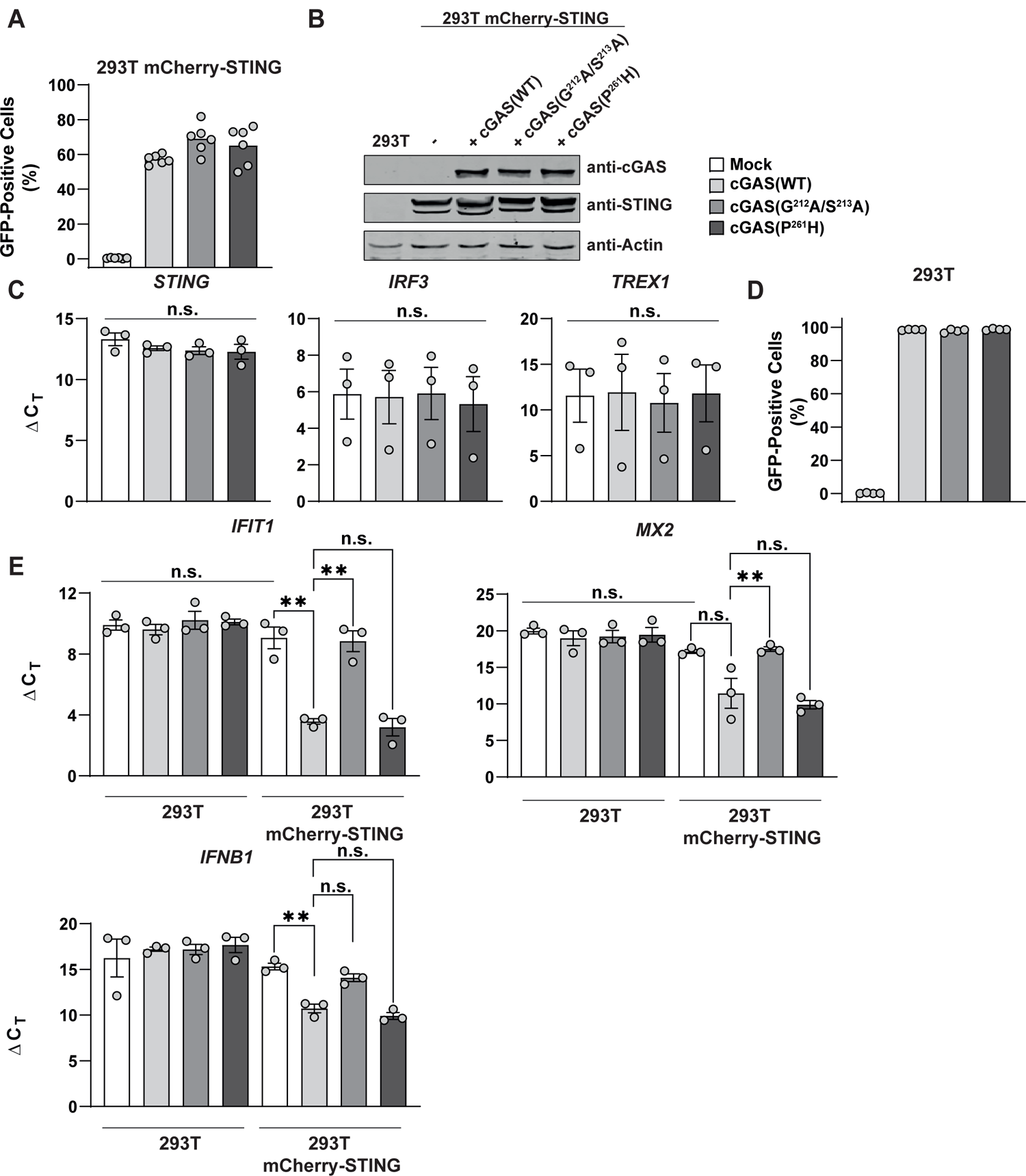
HEK293T cells reconstituted with cGAS and STING display a cGAS-specific ISG expression profile (A) HEK293T cells stably expressing mCherry-STING were reconstituted with individual cGAS-GFP variants. GFP expression was monitored using Flow Cytometry. (B) HEK293T cells that lack endogenous STING expression were reconstituted with cGAS-GFP variants and GFP expression was quantified using Flow Cytometry. (C) Immunoblot analysis of indicated HEK293T cell lysates. One representative blot of two is shown. (D) Expression of *STING*, *IRF3* and *TREX1* mRNA in indicated HEK293T mCherry-STING cells was quantified by RT-Q-PCR and normalized to *RNASEP* mRNA expression. (E) Base line mRNA expression of *IFIT1*, *MX2* and *IFNB1* in indicated cells analyzed by RT-Q-PCR and normalized to *RNaseP* mRNA expression. Error bars indicated S.E.M. from ≥ 3 individual experiments.

